# Pooled-parent exome sequencing to prioritise *de novo* variants in genetic disease

**DOI:** 10.1101/601740

**Authors:** Harriet Dashnow, Katrina M. Bell, Zornitza Stark, Tiong Y. Tan, Susan M. White, Alicia Oshlack

## Abstract

In the clinical setting, exome sequencing has become standard-of-care in diagnosing rare genetic disorders, however many patients remain unsolved. Trio sequencing has been demonstrated to produce a higher diagnostic yield than singleton (proband-only) sequencing. Parental sequencing is especially useful when a disease is suspected to be caused by a *de novo* variant in the proband, because parental data provide a strong filter for the majority of variants that are shared by the proband and their parents. However the additional cost of sequencing the parents makes the trio strategy uneconomical for many clinical situations. With two thirds of the sequencing budget being spent on parents, these are funds that could be used to sequence more probands. For this reason many clinics are reluctant to sequence parents.

Here we propose a pooled-parent strategy for exome sequencing of individuals with likely *de novo* disease. In this strategy, DNA from all the parents of a cohort of unrelated probands is pooled together into a single exome capture and sequencing run. Variants called in the proband can then be filtered if they are also found in the parent pool, resulting in a shorter list of prioritised variants. To evaluate the pooled-parent strategy we performed a series of simulations by combining reads from individual exomes to imitate sample pooling. We assessed the recall and false positive rate and investigated the trade-off between pool size and recall rate. We compared the performance of GATK HaplotypeCaller individual and joint calling, and FreeBayes to genotype pooled samples. Finally, we applied a pooled-parent strategy to a set of real unsolved cases and showed that the parent pool is a powerful filter that is complementary to other commonly used variant filters such as population variant frequencies.

## Background

*De novo* Mendelian diseases are single-gene disorders where the causal variant is found in the proband, but not in the somatic tissues of either of their parents. Such conditions are usually dominant, as the probability of two mutations affecting the same gene is low. *De novo* variants have been identified as the cause of monogenic disorders such as congenital heart disease [1,2], deafness, metabolic disease and a range of syndromic disorders (reviewed in [3]). *De novo* variants are rare, occuring at a rate of ~1.1×10^−8^ per position, or approximately 70 new mutations in each diploid human genome [4]. Yet, in a meta-analysis of diagnostic next-generation sequencing in children, *de novo* variants accounted for the majority of genetic diagnoses in non-consanguineous families [5]. It has also been shown that *de novo* variants are a major cause of neurodevelopmental disorders in non-consanguineous populations [6–9]. In addition*, de novo* mutations are also recognised as contributing to a number of complex conditions such as intellectual disability, autism-spectrum disorders and schizophrenia [10]. Finding the causal genetic variant of a disease can provide diagnosis, prognosis and guide treatment or management [11], yet conditions caused by *de novo* variants can be difficult to diagnose because there is no family history of that condition.

Exome next-generation sequencing (NGS) has become a key tool to discover disease-causing variants. There are two common strategies to finding a genetic diagnosis with exome sequencing: singleton and trio sequencing. In the singleton strategy only the proband is sequenced, while for trio analysis both the proband and their parents are sequenced. The trio approach is particularly powerful in the context of *de novo* mutations (e.g. [6]) where variants observed in the parents can be used as a filter to prioritise those variants in the proband that are likely to be *de novo*. While the trio approach significantly outperforms the singleton approach in terms of diagnostic rate [12], the trio approach is substantially more costly, as it requires library preparation and sequencing of three individuals rather than one. The advantage of the singleton strategy is that while diagnostic rates may be lower, three times as many affected individuals can be sequenced for the same cost, allowing for increased capacity and so more cases overall to be solved. For example, if we sequence 100 exomes and assume a 22% diagnostic rate for singletons and 33% for trios [12] we would expect to solve 22 of 100 cases vs 11 of 33 trios.

In addition to the cost of exome capture and sequencing, we must consider the cost of variant curation. Rather than a specific fee for service, variant curation is usually a limited resource; that is the analyst may only have time to consider a limited number of variants per patient before they must declare that case unsolved and move on to the next patient. Therefore when diagnosing patients from exome sequencing, a key consideration is the number of variants that need to be assessed in each individual. An average individual exome has 10,000-12,000 non-synonymous variants [13], 120 protein truncating variants, and ~54 variants previously reported as pathogenic (although not necessarily relevant to the given phenotype) [14]. This is clearly too many variants for curators or clinicians to assess, so some prioritisation and filtering strategies are necessary. A common strategy is to filter or prioritise variants by their population frequency based on the assumption that highly penetrant pathogenic variants will be rare or absent in unaffected individuals. Frequencies in datasets such as Exome Variant Server [15], 1000 Genomes [16] and the Genome Aggregation Database (gnomAD, previously known as Exome Aggregation Consortium or ExAC) [14,17] are commonly used. For example a frequency of 0.01 might be considered rare, and a frequency of 0.0005 to be very rare [18]. More detailed variant assessment would then consider known pathogenic variants (e.g. from Clinvar), variant consequence prediction (e.g. VEP [19] Condel [20]), evolutionary conservation and clinically-informed gene lists [21]. Even with all of these filtering and prioritisation tools, typically hundreds of variants still remain to be curated and the role of inheritance information is vital in reducing this list. One reason that the diagnostic rate for trios is often higher is that inheritance information can be used to filter out large numbers of variants. Studies that perform trio or other family sequencing can use an inheritance model to select variants that fit with the expected pattern (e.g. *de novo*, dominant, recessive, sex-linked). Yet as we have described, trio sequencing carries a high cost for a modest increase in diagnostic rate.

Here we propose and evaluate a compromise between the singleton and trio strategies: pooled-parent exome sequencing. In this strategy, probands are still sequenced individually. For a given batch of unrelated probands, we pool all their parental DNA, then perform exome capture and sequencing on the pool. Variants called from this pool can then be used as a powerful filter for prioritising *de novo* variants in the probands. Because the exome capture is a substantial portion of the cost (currently ~60%), this strategy provides a dramatic cost saving over a standard trio, while still allowing the the majority of parent alleles to be filtered out when analysing the probands.

Pooled sequencing strategies have been used successfully for assessing population allele frequencies for genome-wide association studies and other applications, reviewed in [22]. More recently pooling has been used in for rare variants in Mendelian disease. For example a recent study used pools of 12 probands to identify *de novo* causes of neurodevelopmental disorders and so were able to detect relevant likely-pathogenic variants in 28% of cases at a greatly reduced cost [23].

In this study we assess the novel strategy of pooled-parent exome sequencing as a method to filter variants from proband exome sequencing. We assume that we are looking for rare *de novo* variants in the probands and therefore the causative variants will not be found in any of the parents. Thus we can take the list of variants called in the probands and filter out any variants observed in any of the parents. We first performed a series of simulations to assess the feasibility of using pooled parents, and to explore the effect of factors such as the number of parents in the pool and sequencing depth. In addition, we compare the performance of two common variant callers on pooled data: GATK HaplotypeCaller [24] and FreeBayes [25]. Finally, we present exome sequencing analysis of four probands with suspected *de novo* causal variants, and a pool of their eight parents. We assess the utility of using the pooled parents as a filter to prioritise *de novo* variants, and compare this strategy to commonly used variant filters, in particular population allele frequency. We show that the pooled-parent filter is a powerful and complementary filter to other strategies.

## Results

### Simulation set up

In order to test the utility of pooled parents for prioritising *de novo* variants we performed a series of simulations. We selected a set of 111 parents from Simons Simplex Collection that had undergone individual exome sequencing [26]. This particular subset of samples was chosen to be matched for DNA sequencer, read length and exome capture kit sequencing depth (see Methods). Only samples with at least 64X median depth over the capture region were retained in order to both remove low quality samples and to limit the range of depths such that if the two samples with the most extreme depths were combined then their reads would appear in a 40:60 ratio. Any other pair of samples combined would have depth ratios between 40:60 and 50:50. This places the simulation within the range of sample ratios likely to be seen with errors in DNA concentration and volume quantification when pooling. The final set consisted of 111 samples, 52 females and 59 males, with median depths ranging from 64X to 97X (median 78X).

We randomly sampled reads from these individuals in various combinations to simulate pooling, then calculated recall rates by comparing variants called on the original exomes to those called on the simulated pools. Full details of our simulation strategies can be found in Methods. Briefly: we simulated pools of two, four, six, eight and ten individuals, generated by extracting reads from a random subset of the 111 exomes. We simulated two different strategies for sequencing depth; constant depth and additive depth. In the constant depth simulations the overall amount of sequencing per pool was kept constant at twice the average depth of the source bams. For example to generate a pool of four 75X samples, half the reads would be drawn from each individual to produce a pool of 150X depth. Since the total depth remains constant, the number of reads sampled from each individual decreases as the number of individuals in the pool increases. For additive depth the total sequencing depth was proportional to the number of samples, so each sample was sequenced to the same depth no matter the pool size, with the total sequencing depth increasing for larger pools. For example, if a pool contained four individuals each with 75X depth, all the reads from all four individuals would be added together to create a 300X pooled sample.

### Choice of variant caller and sequencing depth

When selecting a variant caller for pooled sequencing data, previous comparisons have primarily considered sensitivity and false positive rate [27,28]. For the purposes of a pooled-parent study design we contend that the most important feature of a variant caller is the recall, that is, the number of variants called in the individual that are also detected in the pool. In this application, recall is the most important metric because it affects the number of variants that are able to be filtered from the probands. In contrast to pooling for the detection of pathogenic variants in the pooled samples, here false positive calls are less important. However, we also assess false positives: those variants called in the pool that are not called in any of the individuals. False positives are only problematic in the very unlikely event that they happen to coincide with the causal variant in the proband. So, while previous papers choose their variant caller based on sensitivity, specificity and false positives, we aim to optimise the recall (sensitivity).

A key issue with calling variants in pooled samples is that most variant callers assume the reads come from a single diploid individual. They expect to observe approximately 0%, 50% or 100% of reads supporting a given allele. In our pooled samples we expect many variants to differ dramatically from these ratios. In addition more than two variants can be present at a single locus. Therefore using a variant caller that supports setting ploidy can be advantageous in pooled samples. Both FreeBayes [25] and GATK UnifiedGenotyper [29,30] have been proposed as good variant callers for pooled sequencing data [23,27,28]. Both provide the ability to set ploidy when calling variants, allowing more than two alleles to be called at each locus, and explicitly modelling the lower read counts expected to support rare alleles in pooled data. However UnifiedGenotyper has since been deprecated in favour of GATK HaplotypeCaller [31]. More recently GATK HaplotypeCaller has introduced the option to set ploidy, with values up to 21 possible before reaching performance limitations [32]. Since this change is relatively recent, we are not aware of any published assessment of HaplotypeCaller on pooled data. In addition, HaplotypeCaller is currently the preferred genotyper for unpooled genomes/exomes and variation references such as gnomAD [14,17]. One advantage of using HaplotypeCaller for calling variants in the pool is that it is already the standard analysis tool for individual exomes, and by using the same variant caller for both the proband and the parent pool we reduce the chance of technical artefacts.

The GATK Best Practices now recommends individual calling of samples using HaplotypeCaller followed by joint calling with GenotypeGVCFs [33]. Although it is possible to set ploidy in conjunction with joint variant calling, this is well beyond the intended use for this tool. As such we experienced errors when using joint calling in conjunction with ploidy of 16 and 20 (i.e. our simulated pools of eight and ten). We therefore performed two different analyses with HaplotypeCaller: 1) diploid joint calling on each pool and the individuals that made up that pool and 2) individual variant calling with ploidy set as appropriate for the pool size, with the individuals genotyped separately in diploid mode (see Methods). In addition we compared the performance of these calling modes with FreeBayes.

We compared variants called from the pool to all variants called on the original exomes and calculated recall and false positive rates. For all analysis scenarios the recall across all variants for a pool of two was greater than 94% (Supplementary Table 1). Looking across all variants, the overall trend was for recall to diminish as the pool size increased (Figure 1A). HapolotypeCaller had greater recall than FreeBayes for all simulated pool sizes. This difference was small for pools of two individuals, becoming dramatic in the larger pools. Overall individual calling with HaplotypeCaller performed slightly better than joint calling, especially for larger pools.

**Figure 1:**
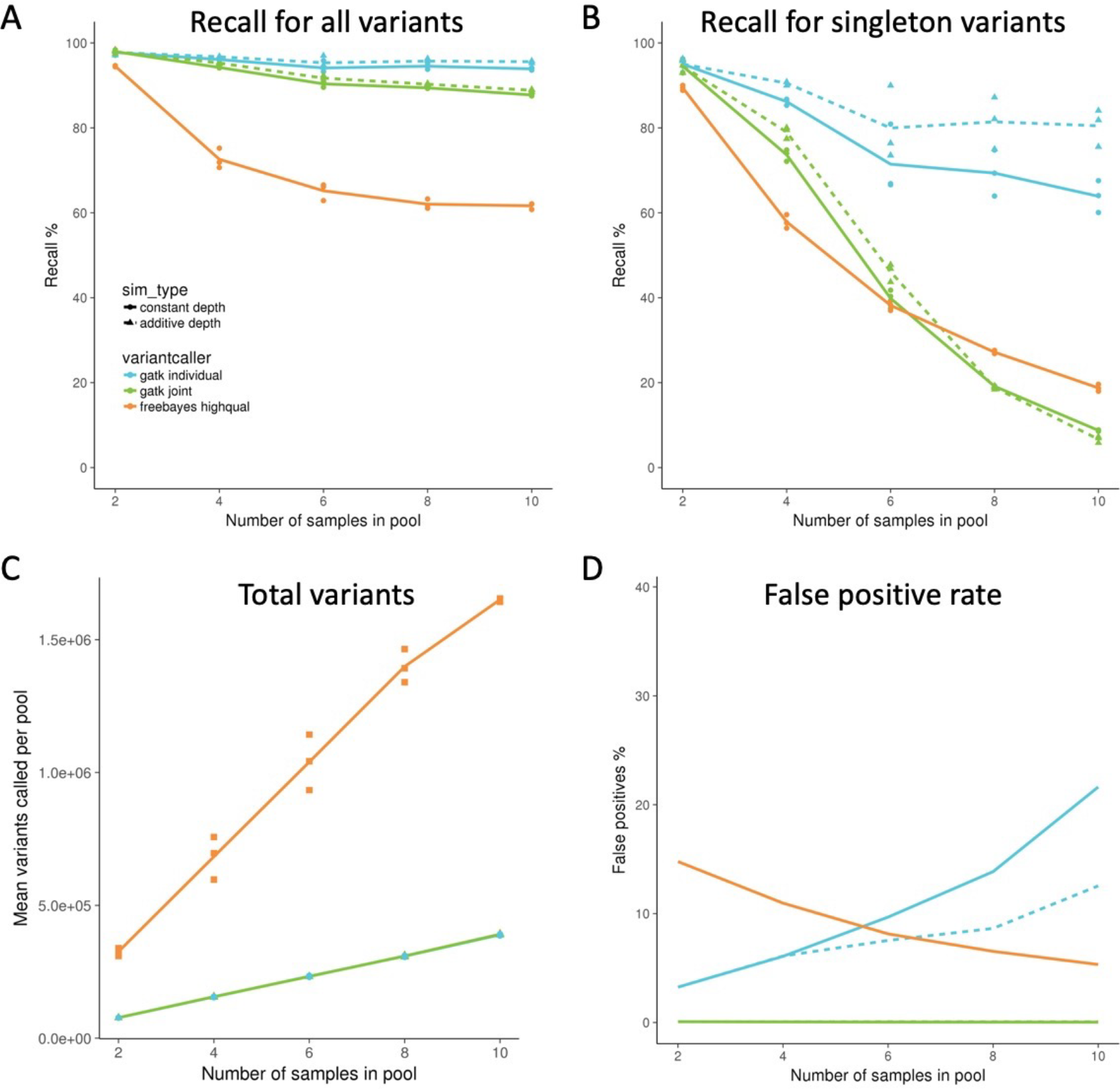
Simulated pools of two, four, six, eight and ten individuals. Each point is a simulation (three replicates) with a different randomly selected set of individuals from a possible 111 individuals. Recall % is the percentage of variants called in all individuals that make up the pool that are also called in the pool. False positive % is the percentage of variants called in the pool that are not called in any of the individuals that make up that pool. A) Recall for all variants B) Recall for variants with an allele present in one copy in one of the individuals C) Total variants called in the pool (for constant depth simulation). HaplotypeCaller individual and joint calling produced similar numbers of variants so are difficult to distinguish at this scale. D) False positive rate for all variants. Mean values for plots A, B, C and D can be found in supplementary tables 1, 2, 3 and 4 respectively.

When using parents to filter for potential de novo variants, the most important and difficult class of variants to recall in the pool are those that are rare and only one allele is likely to be present in the pool. We therefore considered the recall rate for these so called singleton variants. All calling methods had lower recall for singleton variants (Figure 1B). This is expected as these variants will generally have fewer reads supporting them and lower allelic depth. Surprisingly, for HaplotypeCaller, individual calling performed dramatically better than joint calling on these variants. Joint calling showed a pronounced loss of recall with increasing pools size, especially for six or more individuals in a pool. Joint calling draws evidence across samples, thus increasing support for variants found in multiple samples and decreasing support for those variants only found in one sample. This may explain why joint calling performs poorly for singleton variants. It should be noted that FreeBayes reported approximately five to nine times as many variants as HaplotypeCaller, which may contribute to the lower recall rate (Figure 1C, Supplementary Table 3).

For individual calling, HaplotypeCaller saw a steady increase in false positives with pool size (Figure 1D). For joint calling the overall false positive rate is less than 1% and, although it increases slightly with pool size, this increase is insignificant (Supplementary Table 4). So while individual calling with HaplotypeCaller provides superior recall, joint calling better controls the false positive rate. FreeBayes showed a decrease in false positive rate with increasing pool size which is likely why it was previously recommended for variant calling in pooled samples in previous studies.

We further performed an additive depth simulation where all reads from each individual are combined in the pool. In general, increasing the depth increased recall rate and decreased false positive rate. The only exception to this is for singleton joint HaplotypeCaller variants in pools eight and ten. The increase in recall with additive depth was most dramatic for HaplotypeCaller individual calling of singleton variants, where in the largest pool the recall rate increase from 63.9% to 80.5% (Figure 1B, Supplementary Table 2). This indicates that increasing the sequencing depth may be useful in pooled samples. Many of the additive depth pools could not be called using FreeBayes due to massive memory requirements on such large depth samples, so FreeBayes is not included for this simulation.

In summary, HaplotypeCaller individual calling was found to have superior performance in terms of recall rate, especially of singleton variants. In addition, it is more commonly used in clinical variant calling, making this approach more compatible with existing clinical pipelines and reducing the risk of technical bias by using multiple variant callers. Based on our simulations, we expect a pooled parent strategy to provide a useful, cheaper alternative to trio sequencing for *de novo* cases. For example by simply pooling the two parents we get 98% of variants recalled with a low false discovery rate, even with ten in pool we calculated an average recall of 94% (Supplementary Table 1). We found that increasing the sequencing depth can also increase the recall rate.

### Calling variants from real pools

We performed a real pooled-parent sequencing experiment. We had previously exome sequenced four individual probands likely to have *de novo* disease that were still unsolved after the initial variant analysis. We then performed exome sequencing of a pool of all eight of their parents. For the probands, exome capture was performed with the Nextera v1.2 Rapid Exome Capture Kit, and all the libraries were sequenced to ~100× depth over the target region. The parent pool was captured using the Agilent SureSelect QXT Clinical Research Exome kit and sequenced to a median on-target depth of 119× (or ~15× per parent). While the probands and parents were sequenced using different exome captures, the SureSelect Clinical exome is much larger than Nextera, and it mostly covers the same regions so should be able to recover most positions called in the probands. Based on the results of our simulation study we performed variant calling using GATK HaplotypeCaller on each of the samples individually. The probands were genotyped with default (diploid) ploidy and the pool with ploidy set to 16. We then used the parent pool variant calls to filter out variants in the probands. We additionally performed variant annotation with VEP and added gnomAD frequencies (see Methods).

We calculated recall on this real data set as the percentage of variants found in all probands that were also called in the pool. Our calculated recall rate for all variants was 81.3%. This is a little lower than what we expected based on our simulations (Supplementary Figure 1). The recall rate for variants with one allele found in one proband was was 72.6%. This is consistent with the singleton rate observed in simulations (69.4%), however it should be noted that for the real pooled experiment we would expect each proband to have half the genetic material from each parent. Therefore a variant found once in the probands could occur more than once in the parent pool if the variant also appeared in the untransmitted allele. Hence if a variant is found once across all the probands, it could plausibly have an allele frequency of anywhere from 1/16 to 9/16 in the the parent pool and so is not directly comparable with the simulations.

We next used gnomAD to prioritise rare variants by filtering out variants with allele frequency over 0.0005. Of the remaining variants the recall in the pooled sample was 50.8%. This shows while both strategies filter many of the same variants, the parent pool provides substantial filtering beyond gnomAD.

We further assessed the power of using variants called in the parent pool as a strategy for filtering variants in the probands. Here the goal is the minimise the number of variants that need to be curated for each individual by removing those that are unlikely to cause *de novo* disease. We compare using the parent pool as a filter to some of the standard approaches, namely filtering low quality variants and common variants. Figure 2 summarises our variant filtering approach and shows the number of variants remaining after each filter is applied. We defined low quality variant calls as those with QD (QUAL/DP) < 2 [34]. Since our probands have different phenotypes, we expect each to be caused by a different *de novo* variant. So we filtered out variants observed more than twice across the probands, as unlikely to be causal. We also filtered out common variants to retain only very rare variants i.e those observed at a frequency of greater than 0.0005 in gnomAD. We compared this to filtering out variants called in the parent pool. We found that filtering with the parent pool alone resulted in fewer variants than using the gnomAD frequencies alone. Importantly however, we found that the pooled parent filter was a complementary filter to other strategies and reduced the number of variants to less than 45% of the gnomAD only filter (Figure 3A). In particular we note that while gnomAD is useful to filter out variants observed frequently in the general population (specifically those populations included in gnomAD), the pool was able to filter out variants observed in the “private population” made up by these families. This may be particularly important when considering patients from populations that are not well represented in gnomAD. Most of our probands identified as European (Supplementary Table 6), populations which are generally well represented in gnomAD. The gnomAD filter was slightly less effective for the Pacific Islander proband (Proband 3). 95.3% of all raw variants could be filtered using gnomAD for this individual, compared with 96.1-96.4% for the three European probands.

**Figure 2:**
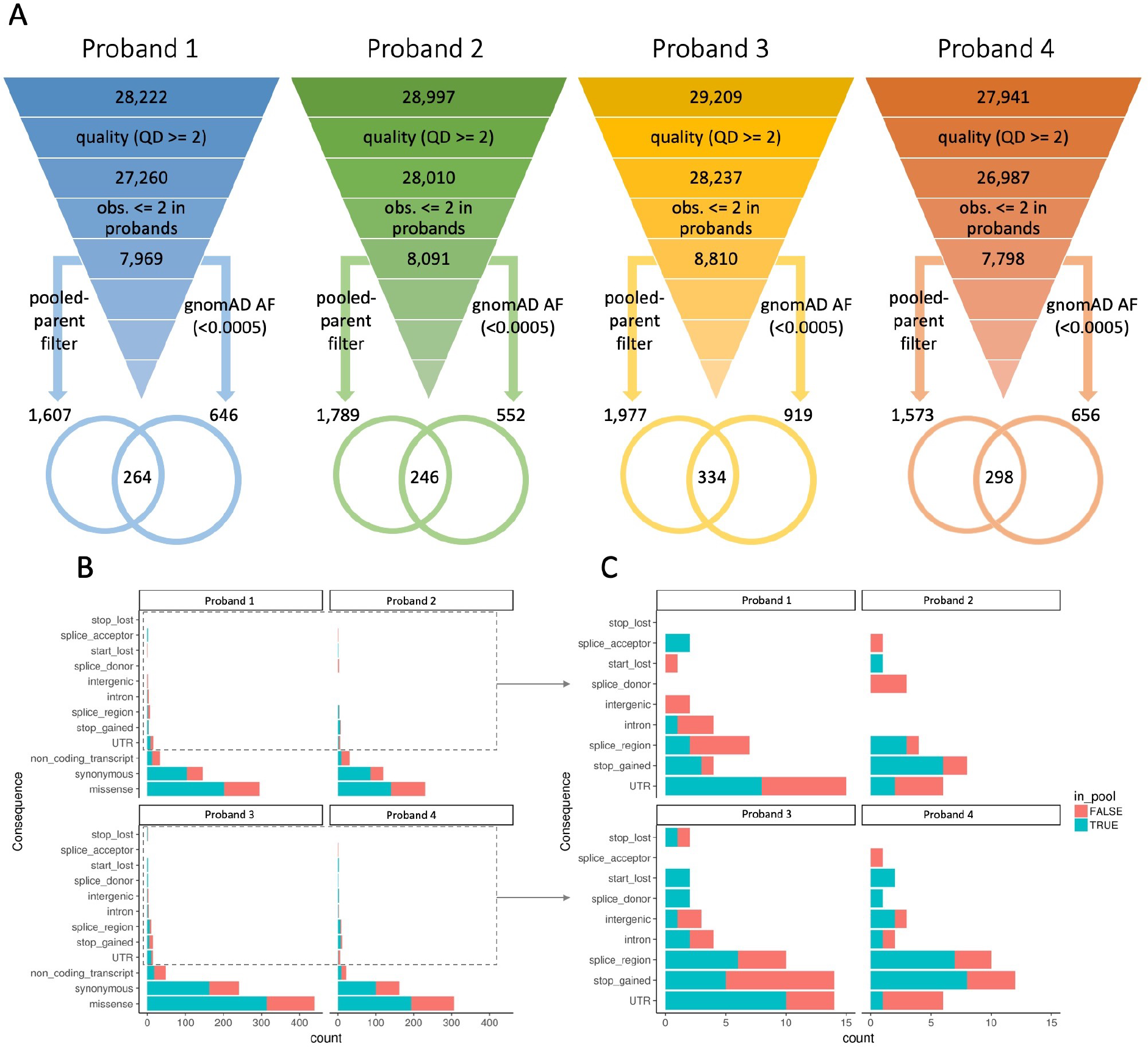
A) Schematic of variant filtering in each proband. Proband IDs are indicated above each inverted triangle. The first figure is the raw number of variants called in each proband. We then filter out low quality variants (QD >= 2) and variants observed 2 or more times across the set of probands, in each case showing the remaining number of variants. At this point variants are filtered with either gnomAD allele frequency or the pooled parent variants, and then the intersection of these two filters is shown as a venn diagram. B) VEP worst consequence annotations for the variants remaining after filtering by quality, frequency in the probands and gnomAD allele frequency. Variants that can be further filtered using the parent pool are indicated as “in_pool”. C) Magnification of lower frequency consequence categories indicated by the dotted rectangles in B.

We also performed annotation with Variant Effect Predictor (VEP) to aid interpretation of potential disease variants. Figures 2B and 2C show the worst consequences annotated by VEP for each variant for only those variants that passed the quality and gnomAD frequency filters. The pooled parent filter dramatically reduces the number of potentially deleterious variants that variant curators might need to consider. For example in all of the probands the parent pool was able to filter out more than half of the missense variants, while in several probands it removed all start lost and splice site donor or acceptor variants.

The pooled parent strategy enabled us to filter out the majority of variants from the probands. Importantly it was complementary, removing a different set of variants to those filtered based on population allele frequency. It removed a number of potentially pathogenic variants, reducing the variant curation load.

## Discussion

The pooled-parent exome sequencing strategy we propose here is a powerful and cost-effective way to prioritise *de novo* variants in the search for causal disease variants. Pooling DNA from parents is dramatically cheaper than full trio exome sequencing. While our simulations indicate that increasing pool size does reduce recall rates, the pooling strategy still allowed filtering of 94-98% of variants (for pool sizes of ten and two respectively). The reduced cost of this strategy allows funds to be reallocated to sequencing of more probands. We assessed variant calling strategies for pooled sequencing, and found that while the GATK HaplotypeCaller joint calling strategy provided the best recall rate overall, HaplotypeCaller individual calling had higher recall for the critical singleton variants. We also found that increasing the sequencing depth for pools was able to increase the recall rate, particularly for singleton variants. Generally increasing the sequencing depth is cheaper than performing additional exome captures. In a real analysis of four probands with undiagnosed likely *de novo* disease we were able to use a pooled-parent strategy to filter over 81% of variants. This strategy was complementary to population allele frequency filtering using gnomAD and resulted in reducing the final list of variants to less than 54% compared to just using gnomAD. Unfortunately, the variants responsible for causing disease in these probands remain unknown.

One reason that the pooled-parent filtering strategy may perform particularly well when compared to population filtering, is that the parent pool is in essence an exquisitely matched population to the probands. The parents are the precise populations from which these probands arose and therefore is an excellent strategy for underrepresented populations. In contrast, gnomAD populations are weighted towards specific populations, particularly individuals of European ancestry [35]. If gnomAD is not a good representation of the population from which the proband arose, then the pooled-parent strategy may perform particularly well in comparison to population filtering.

We have seen the cost of DNA sequencing decrease over time, while the cost of exome capture has remained relatively high, both in reagents and because it is a labour-intensive task. So for exome sequencing the pooled-parent strategy is actually becoming increasingly cost-effective over time. However as the cost of sequencing drops still further, clinicians may increasingly move to whole genome sequencing instead. For whole genome sequencing, the per-sample preparation is a relatively small proportion of the overall cost, so the economics of pooled-parent sequencing are not as compelling. Therefore we expect the pooled-parent strategy to be most useful for exome and other targeted sequencing strategies such as smaller gene panels.

One limitation of this study is the simulations are performed on randomly selected individuals rather than trios. This does not truly reflect the pooling of parents, but rather a comparison of individual samples with those same samples pooled together. However, having the individuals and the pools contain the same samples is a key advantage because the true allele frequencies of variants are known and this design allows false positives to be called. This was particularly useful when assessing the impact of increasing ploidy on the quality of variant calls and in evaluating different variant calling strategies.

## Conclusions

Pooled-parent sequencing is a powerful strategy to filter out inherited variants to allow the analysis to focus on possible *de novo* variants. It is dramatically cheaper than full trio sequencing, allowing additional budget to sequence more probands. Importantly, our analysis shows the pooled parent variant filter is complementary to other standard approaches, in particular, filtering out different variants to using gnomAD population frequencies.

## Methods

All code used for the simulations and analysis of these data sets can be found at https://github.com/Oshlack/pooled-parents-paper.

### Simulating pools

To simulate pooled exome sequencing experiments of various sizes, we combined reads from a set of separately sequenced individuals. We selected parents from the Simons Simplex Collection [26]. These samples were chosen as the largest subset of this collection that were relatively technically homogeneous individuals within the publicly available sequencing data from this project. Specifically this set is matched for DNA sequencer (Illumina GAIIx), read configuration (74 bp paired end reads) and exome capture kit (Nimblegen EZ Exome V2.0). We also removed samples with less than 64× median depth over the capture region in order to both remove low quality samples and to limit the range of sample depths such that min.depth/(min.depth+max.depth) = 0.4. I.e. if any two samples were combined then their reads would appear in a ratio between 40:60 and 50:50. The final set used for simulation consisted of 111 samples, 52 females and 59 males, with median depths ranging from 64 to 97 (median 78). SRA run IDs are listed in the Supplementary Table 2.

Pipelines were written in Bpipe [36]. Raw fastq reads were downloaded for each of these samples, then aligned to gatk.ucsc.hg19.fasta with BWA MEM version 0.7.17 [37] and indexed with Samtools version 1.8 [38] (script: genotype_individuals.groovy).

In the constant depth simulation strategy, pools of two, four, six, eight and ten were simulated by selecting individuals at random from the 111 above, then randomly sampling a proportion of reads from the raw (not deduped) BAM files using samtools view -s (scripts: pooled_sim_bpipe.groovy and pooled_sim_joint.groovy). The proportions of reads were chosen such that the resulting pool would have twice the average depth of the source bams. The sampled bam files are merged with MergeSamFiles (Picard Tools version 2.18.11 [39]), then the read group and sample information from the original samples are removed using Picard AddOrReplaceReadGroups so that – for downstream processes - the BAM will appear to have originated from a single sample. These BAM files were deduplicated using Picard MarkDuplicates.

In the additive depth simulation strategy, pools were simulated so that the total sequencing depth for the pool is proportional to the number of samples. This was achieved by simply combining all reads for the samples in each pool. The simulation steps (scripts: pooled_depth_sim_bpipe.groovy and pooled_depth_joint.groovy) were the same as for the first simulation with the exception that instead of sampling reads from the individual BAMs, all randomly selected BAMs where merged together.

For both the constant and additive simulations we performed three replicates of each pool size using different random seeds and therefore different input samples, resulting in a total of 30 bam files. The code for generating all these simulations can be found at https://github.com/Oshlack/pooled-parents-paper.

### Variant calling and analysis

All variant calling was performed against hg19 and after deduplication with Picard Tools as above. For individual variant calling with GATK HaplotypeCaller version 4.0.10.1 [24], all individual samples were genotyped with default diploid ploidy, while for the pools, ploidy was set to two times the number of samples in the pool. Version 138 of dbSNP was used for all HaplotypeCaller commands. For joint calling, variants were called with GATK HaplotypeCaller -ERC GVCF to generate GVCFs, with default ploidy. Each pool GVCF was combined with the GVCFs of all the individual samples used to create that pool using GATK CombineGVCFs. Joint calling on the pool and its constituent samples was performed with GATK GenotypeGVCFs, to produce a final multi-sample VCF with genotype calls for loci that were called as variant in the pool or any of its individuals. FreeBayes version 1.2.0 [25] variant calling was performed on the individuals and pools from the constant simulation strategy only, as the high depth samples from the additive simulation caused excessive memory consumption. Individual calling was performed with default settings (script: freebayes.groovy), while for pooled variant calling (script: freebayes_pool.groovy) we set the relevant ploidy and ran in pooled-discrete mode with use-best-n-alleles = 4. FreeBayes reports all potential variants, including many of questionable quality, so in both the individuals and the pool we implemented the recommended QUAL > 20 filter [40] using the vcffilter script included with FreeBayes.

To assess recall and false positive rates we compared variants called in each pool to the all variants called on the individuals that made up that pool. For HaplotypeCaller individual calling and FreeBayes we matched up specific variants across VCF files by creating a unique string representing the position and reference/alternate alleles. To do this we created a variant identifier: CHROM_POS_REF_ALT (or ALT1/ALT2 etc if multi-allelic and use this to uniquely match variants across VCF files (filter_individualVCF.py). For the joint calling the pool and individuals were already represented at the same locus in a single VCF file, so we compared a variant across samples in the same vcf (filter_multiVCF.py). The recall rate is then calculated per pool as the number of alleles recalled in the pool divided by the total number of non-reference variants called in all individuals that made up that pool. If an allele was called in the pool but not in any of the individuals we consider it to be a false positive. False positives rate is then the number of false positives called in the pool divided by the total number of non-reference alleles called in that pool. VCF parsing was accomplished using Python 3.6.8 and PyVCF version 0.6.8 [41] and further analysis and plots were generated using dplyr and ggplot2 in R version 3.5.0. Manipulation of exome capture target bed files was performed with Bedtools v2.27.1 [42].

### Samples and sequencing

Four unsolved probands were selected from the Melbourne Genomics Health Alliance Childhood Syndromes project [43] as being good candidates for dominant *de novo* disease based on clinical assessment. This project was approved under Human Research Ethics Committee approval 13/MH/326. Parents provided written informed consent after genetic counseling regarding the testing. As part of the demonstration project these patients received exome sequencing alongside standard of care. DNA was extracted from peripheral blood, and exome sequenced used Nextera v1.2 Rapid Exome Capture Kit on a HiSeq 2500 at the Australian Genome Research Facility to 100X to a median on-target depth of 100×. We additionally sequenced a pool of all eight of these probands’ parents using the Agilent SureSelect QXT Clinical Research Exome capture kit, and 151 bp paired end reads on an Illumina HiSeq4000 to a total median on-target depth of 119×, or on average ~15× per parent.

### Analysis of real pools

The probands and pool were analysed using a similar strategy to the simulations above (scripts: genotype_individuals.groovy and pooled_joint_analysis.groovy). Reads were aligned to gatk.ucsc.hg19.fasta with BWA MEM, indexed and deduplicated. Variant calling was performed with GATK HaplotypeCaller. Each sample was genotyped individually: the probands with default ploidy, and the pool with ploidy = 16. Proband variants were annotated with allele frequencies from gnomAD version r2.0.2 [14,17] using vcfanno version 0.2.9 [44]. VEP was also used to annotate the most severe consequence for each variant.

As for the simulations, we calculated recall as the number of alleles called in any proband that were also called in the pool over all genomic regions (script: filter_individualVCF.py). Multiallelic variants were split to allow individual annotation with gnomAD allele frequencies and VEP consequences. We performed a series of filtering steps. We performed light filtering for variant quality, by filtering out variants with QD < 2 (QD = QUAL/DP), as recommended by the GATK documentation [34]. We also removed variants observed in more than one proband, as they have different diseases so these shared variants are unlikely to be causal. We removed variants with an allele frequency of more than 0.0005 in gnomAD. Finally, we filtered out any variants called in the parent pool. Before examining any individual variants in detail we excluded a set of genes known to cause high penetrance early onset disease to avoid secondary findings in line with Melbourne Genomics ethics requirements (Supplementary Table 8).

## Declarations

### Acknowledgements

We are grateful to all of the families at the participating Simons Simplex Collection (SSC) sites, as well as the principal investigators (A. Beaudet, R. Bernier, J. Constantino, E. Cook, E. Fombonne, D. Geschwind, R. Goin-Kochel, E. Hanson, D. Grice, A. Klin, D. Ledbetter, C. Lord, C. Martin, D. Martin, R. Maxim, J. Miles, O. Ousley, K. Pelphrey, B. Peterson, J. Piggot, C. Saulnier, M. State, W. Stone, J. Sutcliffe, C. Walsh, Z. Warren, E. Wijsman). We appreciate obtaining access to exome sequencing data on SFARI Base (https://base.sfari.org). We thank the patients and families who participated in the Melbourne Genomics Health Alliance study. We also acknowledge the Melbourne Genomics Health Alliance Steering Group, and the Clinical Genomics and Bioinformatics Advisory Groups. We greatly appreciate comments and suggestions on the analysis from Simon Sadedin and Cas Simons. We thank Stefanie Eggers for generating the pooled sequencing data and Andrew Sinclair for contributing to laboratory costs.

### Availability of data and materials

All code used for simulation and the analysis of these data sets can be found at https://github.com/Oshlack/pooled-parents-paper. Approved researchers can obtain the SSC population dataset described in this study by applying at https://base.sfari.org. Individual sample IDs are listed in the Supplementary Table 7. Melbourne Genomics Alliance members and their approved affiliate investigators can apply for access to the real proband data at https://dash.melbournegenomics.org.au/.

### Authors’ contributions

HD and AO conceived the study and drafted the manuscript. HD and KB performed data analysis. ZS, TYT and SMW clinically evaluated and obtained consent from research participants and provided data. All authors read and approved the final manuscript.

### Ethics approval and consent to participate

This study received Human Research Ethics Committee approval (13/MH/326). Parents of patients provided written informed consent after genetic counseling regarding the testing. All individuals have given written informed consent for publication. The experimental methods in this study comply with the Helsinki Declaration.

