## Supplementary Figures and Tables for "Pooled-parent exome sequencing to prioritise *de novo* variants in genetic disease"

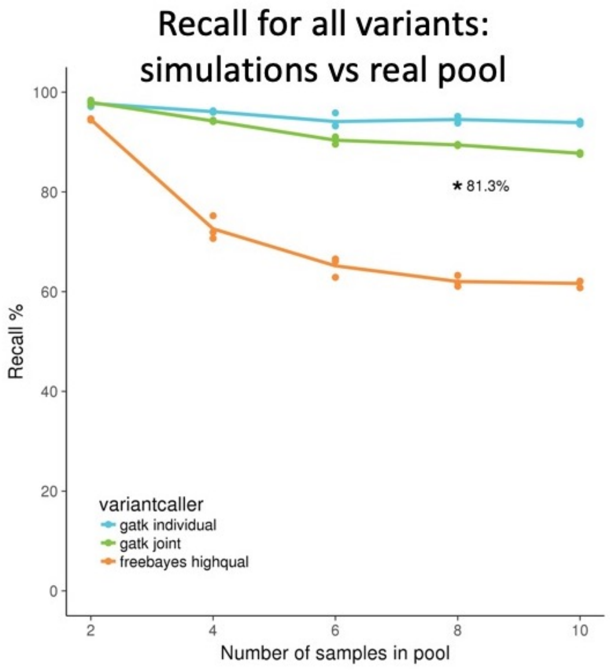


**Supplementary Figure 1: Recall for all constant depth simulations from Figure 1, with * to indicate recall for all variants in the real pools (percentage of variants found in all probands that were also detected in the pool).**

**Supplementary Table 1: Mean recall rate (over three replicates) for all variants as a percentage in simulated pools of two, four, six, eight and ten individuals.** Recall % is the percentage of variants called in all individuals that make up the pool that are also called in the pool.

| | | **Number of samples in pool** | | | | |
|---|---|---|---|---|---|---|
| **sim_type** | **variantcaller** | **2** | **4** | **6** | **8** | **10** |
| **constant depth** | **gatk individual** | 97.8 | 96.1 | 94.1 | 94.5 | 93.9 |
| | **gatk joint** | 98 | 94.2 | 90.4 | 89.4 | 87.7 |
| | **freebayes highqual** | 94.5 | 72.6 | 65.2 | 62 | 61.6 |
| **additive depth** | **gatk individual** | 97.8 | 96.7 | 95.3 | 95.7 | 95.6 |
| | **gatk joint** | 98 | 95.3 | 91.7 | 90.3 | 88.8 |

**Supplementary Table 2: Mean recall rate (over three replicates) for singleton variants as a percentage in simulated pools of two, four, six, eight and ten individuals.** Recall % is the percentage of variants called in all individuals that make up the pool that are also called in the pool.

| | | Number of samples in pool | | | | |
|---|---|---|---|---|---|---|
| **sim_type** | **variantcaller** | **2** | **4** | **6** | **8** | **10** |
| **constant depth** | **gatk individual** | 95 | 86.2 | 71.5 | 69.4 | 63.9 |
| | **gatk joint** | 94.5 | 73.6 | 39.9 | 19.1 | 8.7 |
| | **freebayes highqual** | 89.4 | 57.9 | 38.2 | 27.2 | 18.8 |
| **additive depth** | **gatk individual** | 95 | 90.5 | 80 | 81.4 | 80.5 |
| | **gatk joint** | 94.5 | 79 | 46.1 | 18.8 | 6.7 |

**Supplementary Table 3: Mean variants called per pool (over three replicates) in simulated pools of two, four, six, eight and ten individuals.**

| | | Number of samples in pool | | | | |
|---|---|---|---|---|---|---|
| **sim_type** | **variantcaller** | **2** | **4** | **6** | **8** | **10** |
| **constant depth** | **gatk individual** | 54699 | 74020 | 90934 | 102604 | 124668 |
| | **gatk joint** | 52627 | 68560 | 78770 | 85135 | 93724 |
| | **freebayes highqual** | 310388 | 498533 | 655845 | 790594 | 825890 |
| **additive depth** | **gatk individual** | 54699 | 74040 | 88815 | 96747 | 111552 |
| | **gatk joint** | 52627 | 68553 | 78762 | 85128 | 93708 |

**Supplementary Table 4: Mean false positive rate (over three replicates) as a percentage in simulated pools of two, four, six, eight and ten individuals.** False positive % is the percentage of variants called in the pool that are not called in any of the individuals that make up that pool.

| | | Number of samples in pool | | | | |
|---|---|---|---|---|---|---|
| **sim_type** | **variantcaller** | **2** | **4** | **6** | **8** | **10** |
| **constant depth** | **gatk individual** | 3.25 | 6.08 | 9.68 | 13.9 | 21.6 |
| | **gatk joint** | 0.0735 | 0.0545 | 0.0385 | 0.0349 | 0.0299 |
| | **freebayes highqual** | 14.8 | 11 | 8.14 | 6.53 | 5.33 |
| **additive depth** | **gatk individual** | 3.25 | 6.1 | 7.54 | 8.64 | 12.5 |
| | **gatk joint** | 0.0735 | 0.0643 | 0.066 | 0.0606 | 0.0598 |

**Supplementary Table 5: Mean variants called per individual (diploid) sample used in the simulations for each variant caller.**

| Variantcaller | | |
|---|---|---|
| **gatk individual** | **gatk joint** | **freebayes highqual** |
| 38934 | 39113 | 165067 |

**Supplementary Table 6: Ethnicity of probands, as identified by the patient's family.**

| **Proband** | **Ethnicity** |
|---|---|
| Proband 1 | European (Greek) |
| Proband 2 | Caucasian (Anglo) |
| Proband 3 | Pacific Islander |
| Proband 4 | European (Greek) |

**Supplementary Table 7: SRA run IDs for the 111 exome samples used in pooling simulations. All samples are parents from the Simons Simplex Collection.**

| | | | | | |
|---|---|---|---|---|---|
| SRR1272235 | SRR1301418 | SRR1301506 | SRR1301624 | SRR1301797 | SRR1301865 |
| SRR1272236 | SRR1301419 | SRR1301517 | SRR1301627 | SRR1301801 | SRR1301872 |
| SRR1272246 | SRR1301422 | SRR1301530 | SRR1301631 | SRR1301802 | SRR1301873 |
| SRR1272259 | SRR1301423 | SRR1301533 | SRR1301632 | SRR1301805 | SRR1301876 |
| SRR1272260 | SRR1301430 | SRR1301541 | SRR1301653 | SRR1301806 | SRR1301877 |
| SRR1272263 | SRR1301431 | SRR1301542 | SRR1301654 | SRR1301810 | SRR1301887 |
| SRR1272293 | SRR1301438 | SRR1301545 | SRR1301665 | SRR1301821 | SRR1301888 |
| SRR1272294 | SRR1301439 | SRR1301546 | SRR1301698 | SRR1301825 | SRR1301894 |
| SRR1301242 | SRR1301457 | SRR1301555 | SRR1301699 | SRR1301833 | SRR1301898 |
| SRR1301243 | SRR1301458 | SRR1301556 | SRR1301718 | SRR1301834 | SRR1301899 |
| SRR1301266 | SRR1301461 | SRR1301559 | SRR1301726 | SRR1301841 | SRR1301902 |
| SRR1301267 | SRR1301462 | SRR1301560 | SRR1301737 | SRR1301842 | SRR1301903 |
| SRR1301299 | SRR1301468 | SRR1301567 | SRR1301749 | SRR1301849 | SRR1301918 |
| SRR1301338 | SRR1301469 | SRR1301568 | SRR1301763 | SRR1301850 | SRR1301919 |
| SRR1301339 | SRR1301487 | SRR1301615 | SRR1301764 | SRR1301856 | SRR1515931 |
| SRR1301374 | SRR1301488 | SRR1301616 | SRR1301771 | SRR1301857 | SRR1515932 |
| SRR1301375 | SRR1301491 | SRR1301619 | SRR1301772 | SRR1301860 | |
| SRR1301403 | SRR1301492 | SRR1301620 | SRR1301775 | SRR1301861 | |
| SRR1301404 | SRR1301505 | SRR1301623 | SRR1301776 | SRR1301864 | |

**Supplementary Table 8: List of disease genes excluded from the real analysis (see Methods)**

| | | | | | |
|---|---|---|---|---|---|
| PSEN2 | ALS2 | C9orf72 | SPG20 | FA2H | VAPB |
| PARK7 | CHMP2B | VCP | ATP7B | MAPT | PANK2 |
| TARDBP | UCHL1 | SETX | ANG | GRN | PRNP |
| GBA | SNCA | VPS13A | PSEN1 | NPC1 | |
| ATP13A2 | SNCAIP | OPTN | NPC2 | APOE | |
| PINK1 | PARK2 | TH | SPG11 | FTL | |
| HTRA2 | FIG4 | LRRK2 | FUS | NOTCH3 | |
| UBQLN2 | XK | PLA2G6 | SOD1 | APP | |
